# Field-Programmable Gate Array-Based Ultra-Low Power Discrete Fourier Transforms for Closed-Loop Neural Sensing

**DOI:** 10.1101/2025.02.13.637868

**Authors:** Richard Yang, Heather D. Orser, Kip A. Ludwig, Brandon S. Coventry

## Abstract

Digital implementations of discrete Fourier transforms (DFT) are a mainstay in feature assessment of recorded biopotentials, particularly in the quantification of biomarkers of neurological disease state for adaptive deep brain stimulation. Fast Fourier transform (FFT) algorithms and architectures present a substantial power demand from onboard batteries in implantable medical devices, necessitating the development of ultra-low power Fourier transform methods in resource-constrained environments. Numerous FFT architectures aim to optimize power and resource demand through computational efficiency; however, prioritizing the reduction of logic complexity at the cost of additional computations can be equally or more effective. This paper introduces a minimal-architecture single-delay feedback discrete Fourier transform (mSDF-DFT) for use in ultra-low-power field programmable gate array applications and shows energy and power improvements over state-of-the-art low-power DFT and FFT methods. In a neural sensing application, we observe a 33% reduction in dynamic power and 4% reduction in resource utilization when compared to state-of-the-art FFT algorithms; 38% reduction in dynamic power and 4% reduction in resource utilization when compared to Goertzel Algorithm. While designed for use in closed-loop deep brain stimulation and medical device implementations, the mSDF-DFT is also easily extendable to any ultra-low-power embedded application.

## I. Introduction

**T**HE discrete Fourier transform (DFT) is a fundamental tool in embedded digital signal processing, facilitating the decomposition of input signals into their constituent frequency components. Its utility spans a vast number of applications, making it indispensable for embedded data processing across numerous application domains. Advancement in DFT applications have traditionally focused on reducing computation time and enhancing signal throughput through optimization of numerical algorithms and the hardware on which DFTs are employed. The Fast Fourier Transform (FFT), rooted in the Cooley-Tukey algorithm[1], facilitates rapid computation of DFTs, reducing its computational complexity from 𝒪(*N*^2^) to 𝒪(*Nlog*(*N*)). The FFT reduces time complexity by decomposing calculations into smaller, atomic microoperations which leverages the symmetry and periodicity properties of the DFT to reuse intermediate results and provide temporally efficient calculations[1]. These microoperations are implemented in hardware via complex multiply and accumulate, delay pipeline, and storage ram blocks, commonly referred to as butterfly structures due to their characteristic connections and flow graph[1]–[3]. Extensions of the FFT algorithm have been further optimized to maximize throughput using hardware pipelining techniques or by minimizing implementation size using time-multiplexing[4], [5].

Implantable medical devices are increasingly employing Fourier transform algorithms and processing cores for real-time analysis of biological signaling and control. For example, adaptive deep brain stimulation (DBS), a closed-loop extension of clinical gold standard DBS for the treatment of the motor symptoms of Parkinson’s disease[6] uses measurements of extracellular oscillatory electric potential activity, called local field potentials (LFP), as readouts for motor symptomology and control signals for neural stimulation. In particular, the power in the *β* frequency band (13-30Hz) in the subthalamic nucleus and globus pallidus interna is used as a correlate of bradykinesia. Likewise, increased power in the *γ* frequency band (80-200Hz) is associated with dyskinesia in patients with Parkinson’s Disease[7]–[12]. As closed-loop neuromodulation matures, it is likely that more varied and complex LFP-based biomarkers will be employed for therapeutic sensing, further reinforcing real-time DFT use in implanted devices.

DBS and other electrical stimulation therapies are primarily deployed through implantable pulse generators (IPG)[13]; devices that deliver constant-current electrical stimulation and provide battery management operations. IPGs are chronically implanted and operate from an internal battery and are subject to strict power, hardware, and computational constraints. These constraints place an inherent tradeoff between incorporating advanced features and maintaining device longevity and therapeutic efficacy[14]. Longevity of IPGs is a particularly salient design constraint, with battery replacement necessitating surgical intervention adding potential medical risks, patient stress, and economic costs. Power constraints are even more pronounced in newer generation IPGs which perform biological sensing and signal processing for closed-loop, adaptive neural stimulation at the cost of higher resource and power demands[15] as well as in small animal IPGs[16]– used in preclinical studies, where size and power usage are further constrained by the limited payload capacity of subjects. Efforts to minimize power consumption and optimize the use of limited hardware resources are critical for advancing IPG design. Such advancements could enable the development of smaller, minimally invasive, and potentially injectable devices, broadening the applications of neuromodulation therapies while reducing patient burden.

While the FFT has facilitated fast, high-throughput transforms, the increasingly parallelized architectures required for FFT computation introduce significant dynamic power consumption that directly competes with the power and efficiency requirements for chronic therapeutic stimulation in IPGs. These concerns in other resource constrained settings have been partially addressed through single-delay feedback fast Fourier transform methods (SDF-FFT), a class of memory efficient FFT implementations using a single multiply line with coefficients stored in feedback shift registers[20]–[23]. This class of architecture reduces memory loads over conventional highly-parallelized FFT implementations at the cost of increased latency[22], [23]. While these approaches attempt to minimize RAM and transistor usage, they are still bound by static and dynamic power loads owing to the complex computational structure of the FFT. Additional approaches, such as architectures using approximate multiplication and addition operations[24], can improve energy efficiency but often degrade signal resolution, an unacceptable compromise for applications requiring high-resolution Fourier representations.

Energy-efficient discrete Fourier transform (DFT) computations are commonly achieved by limiting the evaluation of Fourier coefficients to a small, predefined subset of frequency bins. One widely adopted technique is the Goertzel algorithm, which is considered a standard for low-power and embedded applications. This method reformulates the DFT as a series of second-order recursive filters, enabling the efficient computation of real-valued Fourier coefficients for selected frequencies [25], [26]. These target frequencies are typically known *a priori*, allowing for significant reductions in computational complexity. However, a notable limitation of the Goertzel algorithm is its inability to preserve full phase information, which can be critical in certain applications such as adaptive deep brain stimulation (DBS) [27], where both amplitude and phase dynamics may be relevant. Alternatively, burst I/O fast Fourier transforms (FFTs) improve energy efficiency by decomposing the FFT process into discrete operational phases—data loading, coefficient computation, and data unloading. This staged approach reduces peak resource demands but may result in increased overall computation time.

This work presents the design, implementation, and evaluation of a power-efficient minimal single-delay path discrete Fourier transform (mSDF-DFT) architecture for use in ultra-low power embedded applications that require online spectral estimation. The mSDF-DFT is a direct implementation of the DFT that computes in a time-multiplexed fashion. We show that simplified DFT architectures can be more power and resource-efficient than the more computationally efficient FFT architectures by maximally reducing logic complexity while maintaining DFT accuracy and time complexity. The mSDF-DFT shows improvements in power and resource use while providing online performance comparable to embedded standard FFT implementations. Comparisons with state of the art (SOTA) Xilinx burst I/O FFT, pipelined FFT and canonical Goertzel Algorithm[28] are made to characterize and elucidate the advantages of our new mSDF-DFT architecture. Lastly, we validate the use of the mSDF-DFT in neural sensing applications by calculating LFP *β* and *γ* power bands for use in a closed-loop DBS application. We show that, in this application, mSDF-DFT outperforms existing low-power FFT and low-power DFT methods in both resource utilization and dynamic power consumption.

## II. ALGORITHM IMPLEMENTATION

The mSDF-DFT and SOTA benchmark DFT and FFT algorithms were designed in SystemVerilog and evaluated on a Xilinx Spartan-7 FPGA (Boolean Board, Real Digital, Pullman, WA) with synthesis and analysis performed in Vivado Design Suite (AMD). All hardware architectures included runtime configurable (RTC) and non-runtime configurable (Fixed) implementations. RTC variants allowed setting of DFT parameters, such as transform length and number of frequency bins, at runtime at the cost of more complex architecture structures, while fixed implementations set DFT parameters before hardware synthesis. Available RTC parameters for each DFT are given in Table I. Input data, phase factors, and all intermediate results were expressed with 12-bit fixed point data format with 4 fractional bits. Truncation was used for rounding, and all DFT architectures were implemented with an 80 MHz system clock. mSDF-DFT architecture performance was compared against benchmark Goertzel filter, burst I/O FFT, and pipelined FFT architectures representing current standard performance in hardware resource use, runtime and throughput efficiency, and power use in embedded systems, and are described in subsequent sections.

**TABLE I.**
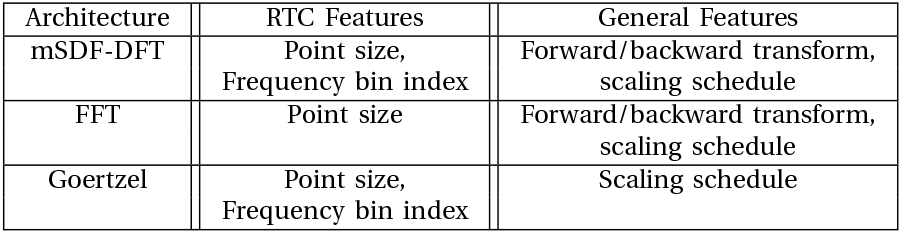
Summary of Architecture Runtime Configurable Features.

| Architecture | RTC Features | General Features |
| --- | --- | --- |
| mSDF-DFT | Point size,<br>Frequency bin index | Forward/backward transform,<br>scaling schedule |
| FFT | Point size | Forward/backward transform,<br>scaling schedule |
| Goertzel | Point size,<br>Frequency bin index | Scaling schedule |

### A. Implementation of the Goertzel Algorithm

The Goertzel filter[28] is a commonly used modification of the DFT which utilizes the periodicity of the 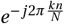 terms to reduce computational loads, using only real-valued coefficients and limited memory to facilitate efficient implementation in embedded applications. Goertzel filter operates on the input *x*[*n*] in two stages. The first stage produces a real-valued intermediate sequence *s*[*n*]:

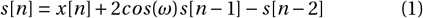

where *ω* is given by 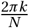, *k* is the frequency bin index, and N is the width of the transform window. The second stage produces the complex output sequence y[n]:

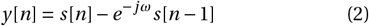

representing a convolutional form of the DFT. Goertzel filter architectures were implemented as an FSM, with one multiplier, one adder, and one ROM unit.

### B. Implementation of the Pipelined FFT

Decimation-in-frequency pipelined FFTs were implemented using the Xilinx FFT IP Core (AMD). The pipelined FFT implementation decomposes the input signal into parallel streams of add/multiply, twiddle storage ROM, and radix butterfly structures, allowing for temporally efficient FFT calculations at the potential cost of hardware complexity[29]. Pipelined architectures were implemented as radix-2 decomposition networks consisting of *log*_2_(*N*) stages, each with 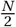 radix-2 butterflies.

### C. Implementation of the Burst I/O FFT

Burst I/O FFT methods represent a tradeoff between low-resource Goertzel implementations and fully parallelized FFTs. Burst I/O architectures decouple DFT computation from data input and output operations, performing serial processing of blocks of input signals, as opposed to pipelined FFTs, which process signals on arrival to Fourier transform cores. Burst I/O architectures provide minimal memory overhead at the cost of increased computational latency. Burst I/O FFTs were implemented as radix-2 employing a shared additive line with a radix-2 butterfly that shares one adder serving to perform parallelized decimation in time computation with minimal memory overhead at the expense of increased computation latency. Comparative FFTs were all implemented in Xilinx Vivado studio using the Xilinx Fast Fourier Transform 9.1 IP core (AMD).

### D. Minimal SDF-DFT Implementation Details

The DFT of an N-point discretized signal *x*_*n*_ for *k* frequency samples is defined as:

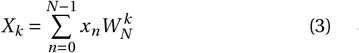

for *k =* 0, 1,… *N −* 1. The 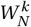 term is the periodic basis defined as 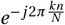 and called the phase or twiddle factor. The mSDF-DFT is implemented as a direct computation of the governing Fourier equation with a finite state machine (FSM) implementation given in Fig. 1A and algorithm 1. In contrast to traditional FFT structures, the mSDF-DFT performs direct computations around a predefined number of frequency bins, creating a computational complexity of 𝒪(*N × N*_*B*_), where *N*_*B*_ is the number of frequency bins to be processed, and N is the point size. The module consists of one multiplier, one adder/subtractor, and one read-only memory (ROM) containing the phase factors (Fig. 1C). The input is a stream of N complex values, each represented by a pair of 12-bit-wide twos-complement numbers. The real and imaginary components of each sample are processed separately with a time-multiplexed approach. The compile-time parameters of the module consist of *N*_*B*_ and maximum point size (*N*_*M AX*_). As such, the mSDF-DFT trades time-complexity for savings in dynamic power and FPGA resource utilization, paramount to ultra-low power embedded processing.

**Fig. 1.**
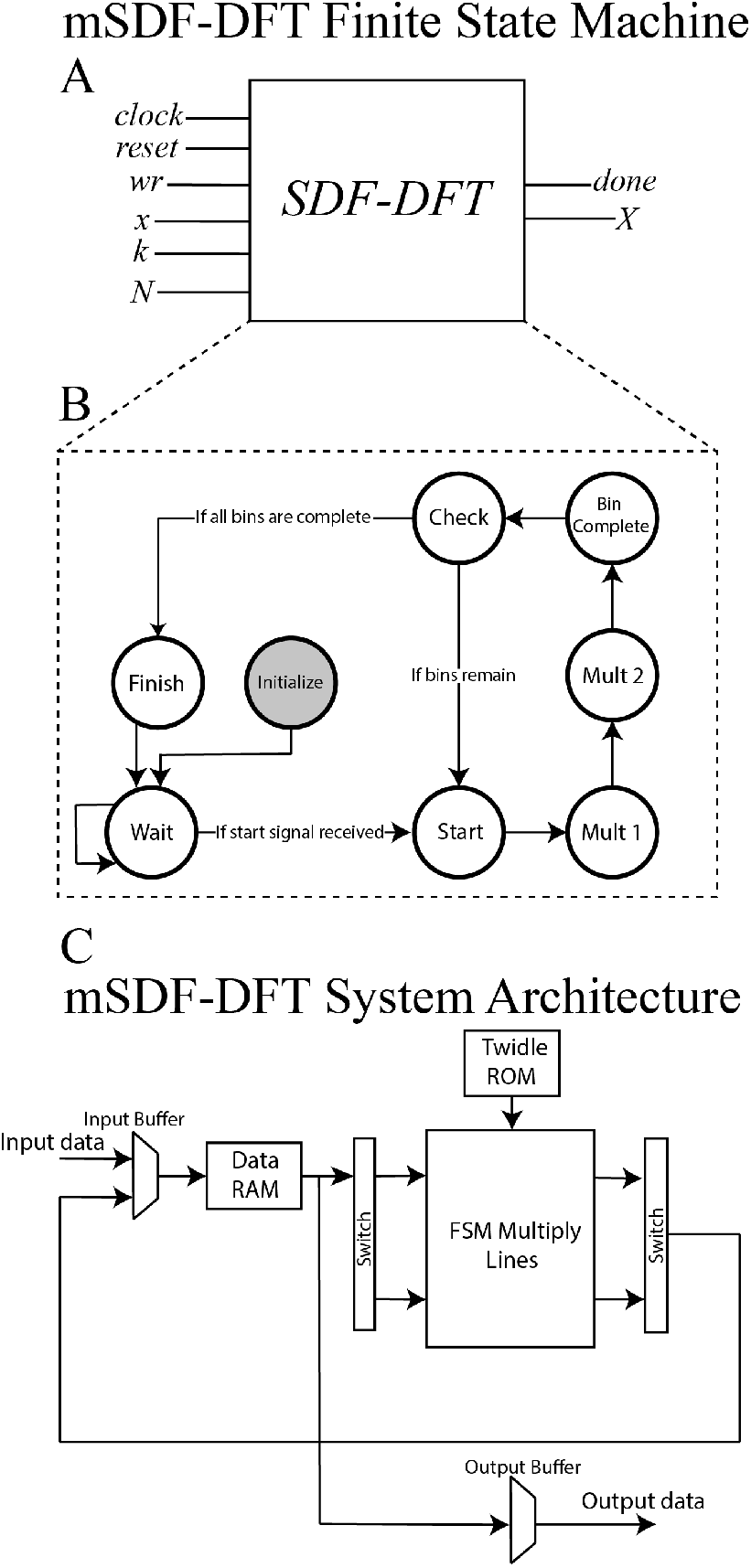
Schematic description of the SDF-DFT finite state machine. A. Inputs to the SDF-DFT include clock, reset, write (wr), input time series (x), frequency bin index (k), and transform window (N). Outputs consist of output FT representation (X) and binary done indicators. B. Flow diagram of the SDF-DFT finite state machine. C. Hardware structure diagram of the mSDF-DFT.

The architecture operates through a finite-state machine (Algorithm 1) comprising eight distinct states (Fig. 1.B). Upon power-up, the architecture transitions into the **Initialize** state, which is executed a single time during system startup. In this state, one-quarter of the complete phase factor table is precomputed and stored in ROM. This optimization significantly reduces ROM usage by exploiting the symmetry and periodicity of the complex exponential function: the remaining three-quarters of the phase factors can be derived from the first quarter via successive multiplications by the complex unit *−j*. The real and imaginary components of the complex exponentials are stored in separate lookup tables referred to as the cosine table (cos) and sine table (sin). For clarity, this subset of phase factors is referred to as the first-quarter phase factors. The real and imaginary components of these complex exponentials are respectively stored in separate lookup tables, subsequently referred to as the cosine table (cos) and sine table (sin). For clarity, this subset of phase factors is henceforth referred to as the first-quarter phase factors. Additionally, during the **Initialize** state, the index of the initial frequency bin to be computed is loaded into the system. Following the **Initialize** state, control transitions to the **Wait** state, in which the first sample is polled with the first-quarter phase factors loaded from ROM by the following operation:

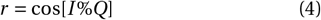

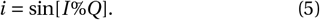

where *r* is the real component of the first-quarter phase factor, and *i* is the corresponding imaginary component. *I* refers to the index of the phase factor in the full phase factor table, % refers to the modulo function, and *Q* is defined as:

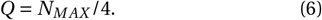

Upon receipt of a new sample, a start signal is asserted and the system transitions into the **Start** state, where the appropriate complex phase factor is reconstructed. This is achieved by selectively negating or retaining the signs of the real (r) and imaginary (i) components of the stored first-quarter phase factor, based on the current frequency bin index. This operation serves to map the index (*I*) to the corresponding location in the phase factor table. Upon completion of the **Start** state, the architecture proceeds through two multiply states, **Mult1** and **Mult2** respectively. Both multiple states share a hardware multiplication unit to conserve hardware resources and reduce power consumption. In the **Mult1** state, the complex input sample is multiplied by the real component of the phase factor, with the resulting computation stored in an accumulation register for further processing. The **Mult2** state then multiplies the input sample by the imaginary component of the phase factor, with the resulting computation stored in the accumulation register. After the multiplication states, the architecture proceeds to the **Check** state, where the architecture increments a sample counter, compares it with the configured *N* value (Fig. 1.A), and checks if all input samples are processed. If more bins remain, the architecture returns to the **Start** state and iterates through all remaining bins. Upon completion, the architecture signals a complete state, sequentially prints out the computed values, and returns to a **Wait** state.

#### Algorithm 1 Minimal SDF-DFT Finite State Machine.

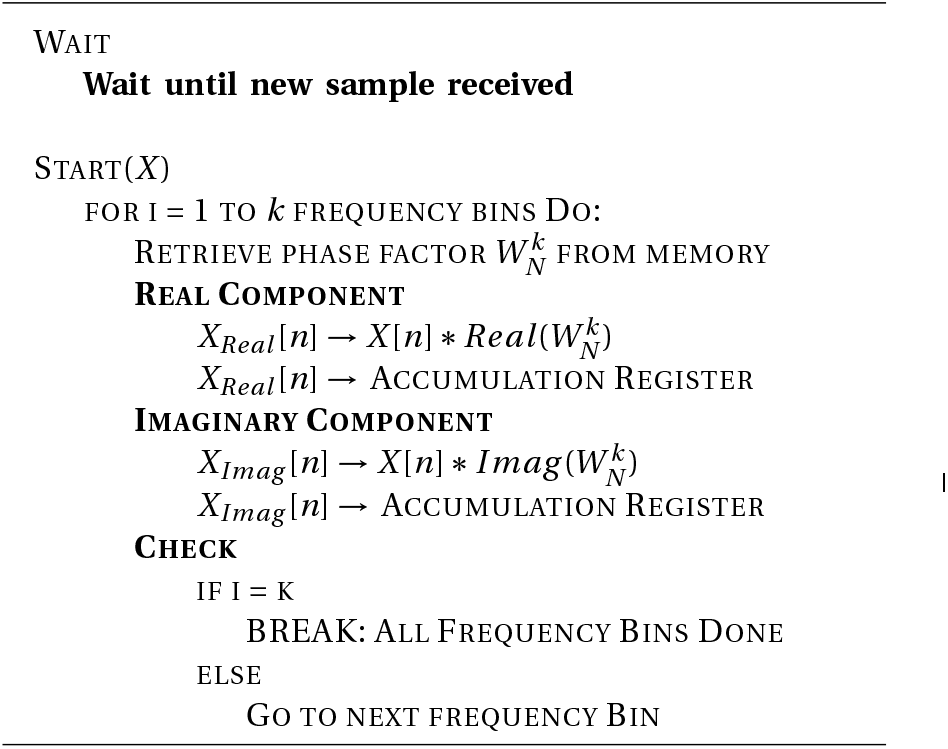

### E. Performance Metrics

Performance evaluations are focused on dynamic power and physical resource utilization. Total system power was calculated using the Vivado power estimation toolset, with total power consumption defined as:

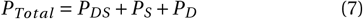

with *P*_*D*_ *S,P*_*S*_, *P*_*D*_ denoting Spartan 7 static power consumption, design architecture static power consumption, and design architecture dynamic power consumption, respectively. Design architecture dynamic power consumption was estimated for forward Fourier transform operations for N-point transforms between 32 and 32,768 points. mSDF-DFT and Goertzel filter architectures contain an additional number of frequency bands parameter (*N*_*B*_) with *N*_*B*_ tested between 4 to 64 for all N point sizes.

To evaluate the total number of physical resources used by each DFT architecture, a physical resource utilization metric was defined as:

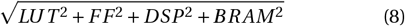

where *LUT, FF, DSP*, and *BRAM* refer to percent utilization measured in Vivado Design Suite of available look-up tables, flip flops, digital signal processing cores, and block random access memory, respectively. Physical resource utilization was estimated for forward Fourier transform operations with N between 32 and 32,768 points and *N*_*B*_ between 4 and 64.

Comparisons between mSDF-DFT and state-of-the-art architecture performance were estimated as:

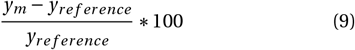

where *y*_*m*_ is the value of physical resource utilization or dynamic power for the mSDF-DFT and *y*_*reference*_ is the physical resource utilization or dynamic power for the comparative benchmark architecture, respectively. The pathological case of calculating DFTs with transform length of 64 with 32 bins (32,64) was excluded from analysis as this set of parameters created an under constrained degenerate Fourier transform.

Architecture latencies were evaluated as the number of clock cycles elapsed between the loading of the initial input sample and the egress of the first computed frequency bin, and the unloading of the first computed frequency bin. Latency of the pipelined and burst I/O transforms implemented with Xilinx IP cores were obtained with Vivado simulations. Least-squares linear regression was performed to obtain a linear estimate of latency as a function of the Fourier transform point size.

## III. EVALUATION OF mSDF-DFTS FOR NEURAL SENSING

To evaluate online neural sensing with the mSDF-DFT method, raw trace recordings from auditory cortex in response to medial geniculate body infrared neural stimulation were used. LFP recordings were used from a previous experiment (study results, stimulation conditions, and ethics approval can be found in[30]). Infrared neural stimulation (INS) is an optical neuromodulation technique that uses coherent infrared light to elicit spatially constrained excitatory neural responses in nerve and neuron[31]–[34] without electrical stimulation artifact.

To mimic common IPG closed-loop sensing, sample LFPs were loaded into the memory of a nRF5340 microprocessor (Nordic Semiconductor, Trondheim, Norway) and transmitted to a Xilinx Spartan 7 FPGA (AMD) through a custom serial peripheral interface (SPI) driver. *β* (13-30 Hz) and *γ* (30-100 Hz) band power are calculated with mSDF-DFT and transmitted back to the microprocessor. Specifications of the neural sensing implementation are summarized in Table 2. Ground truth spectral content of test LFPs was calculated offline in Matlab (Mathworks, Natick MA) using the same number of bins and transform lengths as online mSDF-DFTs. Error between online calculation and ground truth was calculated as:

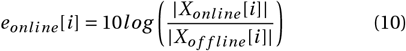

where *X*_*online*_ [*i*] and *X*_*offline*_ [*i*] are the complex amplitude for frequency bin *i* for the online and Matlab calculations, respectively. The application is compiled separately with the four DFT/FFT methods, and their performance is compared.

**TABLE II.**
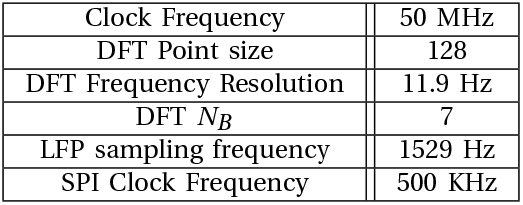
Specification of the Neural Sensing System.

## IV. Results

In the following sections, we first compare the performance of mSDF-DFT across different configurations against SOTA algorithms. Then, we evaluate their latency characteristics as a function of input size. Lastly, we compare the performance of these algorithms in a specific online neural sensing application and show that mSDF-DFT outperforms SOTA algorithms in both resource utilization and dynamic power consumption. Minimal SDF-DFT synthesis and implementation results in Vivado are summarized in Table 3. The mSDF-DFT was implemented with the inverse DFT features to provide appropriate comparisons against the Xilinx IP cores; thus, further reduction in power and resource consumption can be easily realized by removing the feature if only Fourier decomposition is needed for the application.

**TABLE III.** Summary of Selected mSDF-DFT Implementations in Vivado.

| Variant | $N_B$ | $N$ | LUT count | Flip-flop count | DSP count | BRAM count | Dynamic Power (mW) |
| --- | --- | --- | --- | --- | --- | --- | --- |
| RTC | 8 | 512 | 360 | 353 | 2 | 1 | 9 |
| RTC | 16 | 512 | 477 | 546 | 2 | 1 | 10 |
| RTC | 32 | 512 | 954 | 936 | 2 | 1 | 15 |
| Fixed | 8 | 512 | 287 | 342 | 2 | 1 | 8 |
| Fixed | 16 | 512 | 363 | 535 | 2 | 1 | 9 |
| Fixed | 32 | 512 | 902 | 926 | 2 | 1 | 14 |

### A. Performance Evaluation of mSDF-DFT

The mSDF-DFT was designed to provide ultra-low dynamic power consumption and minimal resource usage for a minimal reduction in time-complexity. mSDF-DFT performance was benchmarked against Goertzel filter DFTs as well as parallelized burst I/O and pipelined FFT methods. We found that mSDF-DFT’s performance was consistent across the RTC and the fixed variants with respect to the pipelined and the burst I/O FFT. Comparisons were made in both non-RTC (Figure 2) and RTC (Figure 3) settings to account for differential memory usage for parameters set pre vs post compilation. It was observed that the magnitude of power reduction was greater than the savings in resource usage. The mean resource saving was 14.2% with a standard deviation of 198% while mean power saving was 53.3% with a standard deviation of 43.1%. Minimal SDF-DFT achieved significant reduction in both power and resource consumption relative to the pipelined FFT at all parameters while it outperforms the burst I/O FFT with low *N*_*B*_ to N ratio (Fig 2A,2C,3A,3C). More specifically, this ratio was found to be 0.5 at point size of 32 and diminishes approximately in a power series fashion as a function of point size. This relationship can be modeled by:

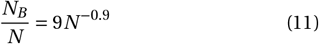

**Fig. 2.**
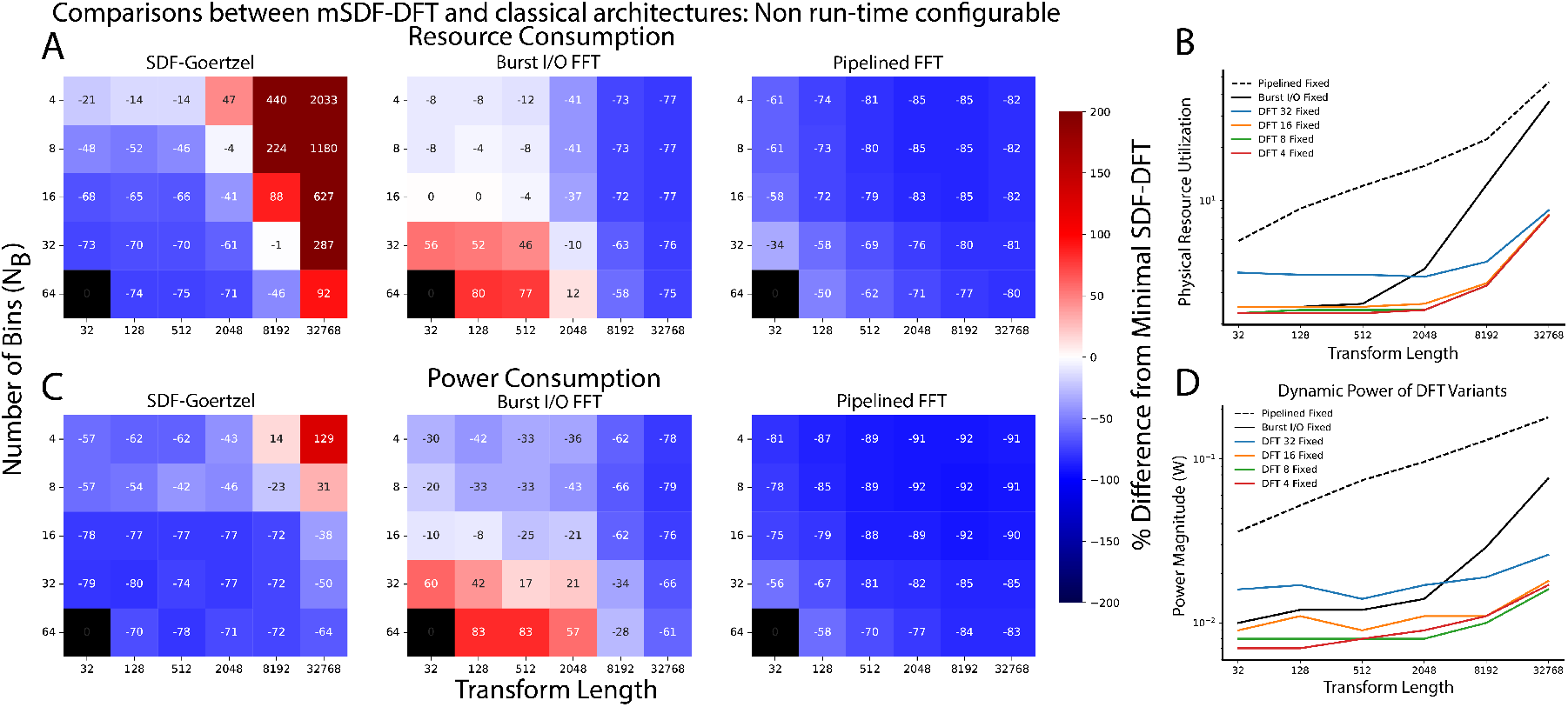
Comparison of non-runtime configurable mSDF-DFT and traditional FFT architectures. A,B. Heatmap of the resource usage (A) and power (B) performance of the mSDF-DFT relative to traditional algorithms. At each combination of (the total number of frequency bins (*N*_*B*_), point size(N)), the percent difference of the mSDF-DFT’s dynamic power and resource consumption relative to the Goertzel, burst I/O FFT, and the pipelined FFT was calculated. The performance values of the FFTs are estimated with *N*_*B*_ equal to N. The combination (*N*_*B*_ =64, Transform length = 32) was manually set to 0 as number of bins exceeds transform length. Blue shades indicate decreased resource and power consumption of mSDF-DFT relative to test algorithms, i.e. better performance. C,D. Line plots of performance for resource (C) and power (D) consumption. These plots represent the same data shown in A,B cast against mSDF-DFT *N*_*B*_ parameters of 4,8,16, and 32.

**Fig. 3.**
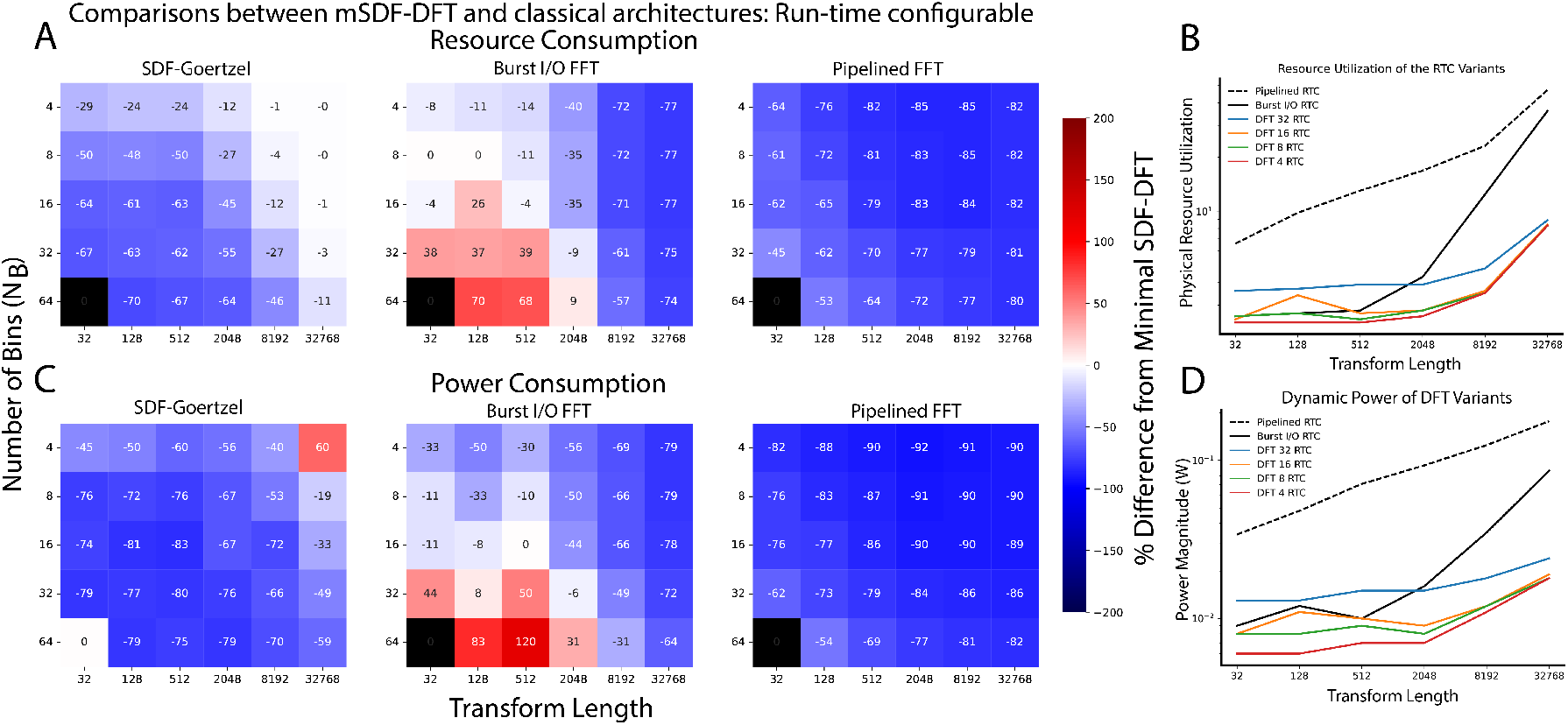
Comparison of runtime configurable mSDF-DFT and traditional FFT architectures. A,B. Heatmap of the resource-usage (A) and power (B) performance of the mSDF-DFT relative to traditional algorithms. At each combination of (the total number frequency bins (*N*_*B*_), point size(N)), the percent difference of the mSDF-DFT’s dynamic power and resource consumption relative to the Goertzel, burst I/O FFT, and the pipelined FFT was calculated. The performance values of the FFTs are estimated with *N+* _*B*_ equal to N. The combination (*N*_*B*_ =64, Transform length =32) was manually set to 0 as number of bins exceeds transform length. Blue shades indicate decreased resource and power consumption of mSDF-DFT relative to test algorithms, i.e. better performance. C,D. Line plots of performance for resource (C) and power (D) consumption. These plots represent the same data shown in A,B cast against mSDF-DFT *N*_*B*_ parameters of 4,8,16, and 32.

Minimal SDF-DFT exhibits clear advantages in both resource use and power consumption over the FFTs at point size greater than 2048, where resource use reduction ranges from 57% to 85%, and power reduction ranges from 31% to 91% (Fig 2B, 2D, 3B, 3D). This significant reduction is mainly attributable to the fact that the mSDF-DFT only calculates 0.01% to 0.8% of the frequency components relative to the FFTs.

Minimal SDF-DFT architectures outperformed Goertzel Filter in both non-RTC (Figure 2A,B) and RTC (Figure 3A,B) conditions except at larger bin sizes and transform lengths in which Goertzel begins to dominate. This is likely due to its efficient BRAM access patterns and storing minimal phase factors when the frequency bin indexes are fixed. However, mSDF-DFT showed clear advantages for small to moderate transform lengths and bin sizes relevant to low power and resource usage implementations. Taken together, this data suggests that *N*_*B*_ and transform sizes can be finely tuned to achieve minimal dynamic power and resource usage to facilitate transforms across a variety of application constraints. This data also suggests that mSDF-DFT is particularly advantageous when desired frequency bands are known *a priori*, such as in adaptive DBS where *β* (13-30 HZ) and *γ* (30-100 HZ) LFP activity provide controllable biomarkers for Parkinson’s disease[35], [36]. It is important to note that, unlike the Goertzal algorithm, the mSDF-DFT provides low power computation of full signal frequency and phase information even when there are no target frequency bands of interest.

### B. mSDF-DFT Latency Evaluation

A fundamental tradeoff between mSDF-DFT and canonical FFT implementations is time complexity vs dynamic power and resource loads. Latency calculations were thus performed to quantify time tradeoffs vs mSDF-DFT *N*_*B*_ against canonical FFT methods. Latency of the mSDF-DFT was found to be *N ×* (5 *×N*_*B*_ *+* 1) and *N ×* (4 *×N*_*B*_ *+* 1) for the RTC and the fixed variants, respectively. For each sample, one clock cycle was required for initiation and 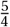 clock cycles were required to process each frequency bin. Similarly, Goertzel filter’s latency was found to be *N* (4*×N*_*B*_ *+*1)*+N*_*B*_ *×*4. The burst I/O FFT featured an approximate latency of 16.8 *× N* while both variants of the pipelined FFT featured an approximate latency of 2 *×N*. The mSDF-DFT displayed greater latency than the FFTs (Fig 4) compared to other FFT methods as expected. Minimal SDF-DFT latency displayed linear growth as a function of transform length across all *N*_*B*_ values. However, we observed that mSDF-DFTs can achieve comparable time performance to burst I/O FFTs with longer transform lengths and smaller *N*_*B*_. Quantification of latency can therefore allow optimal choice of mSDF-DFT parameters to satisfy computational time costs across application. While the mSDF-DFT exhibits a worse asymptotic runtime complexity than FFT methods, mSDF-DFT transform time is dependent on the total number of frequency bins used (𝒪(*N ×N*_*B*_)). Therefore, *a priori* knowledge of the frequency band of interest can significantly reduce algorithm latency. This is often the case with neural sensing applications, where the frequencies of interests are well-defined. For instance, *β* band signals have a period of around 50 ms. A response delivered within 5 ms of signal detection can be considered to be concurrent. Assuming an FPGA clock speed of 50 MHz, 1 kHz sampling rate (*f*_*s*_), and a frame size of 64, the SDF-DFT incurs 0.0064 ms latency, while burst I/O FFT and pipelined FFT yield 0.022 ms and 0.0027 ms, respectively. At a frame size of 1024, mSDF-DFT incurs 0.68 ms latency, while burst I/O FFT and pipelined FFT yield 0.34 ms and 0.04 ms, respectively. Crucially, fewer frequency bins need to be calculated at smaller frame size since the number of frequency bins that span a particular frequency band is:

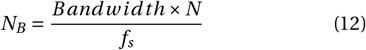

**Fig. 4.**
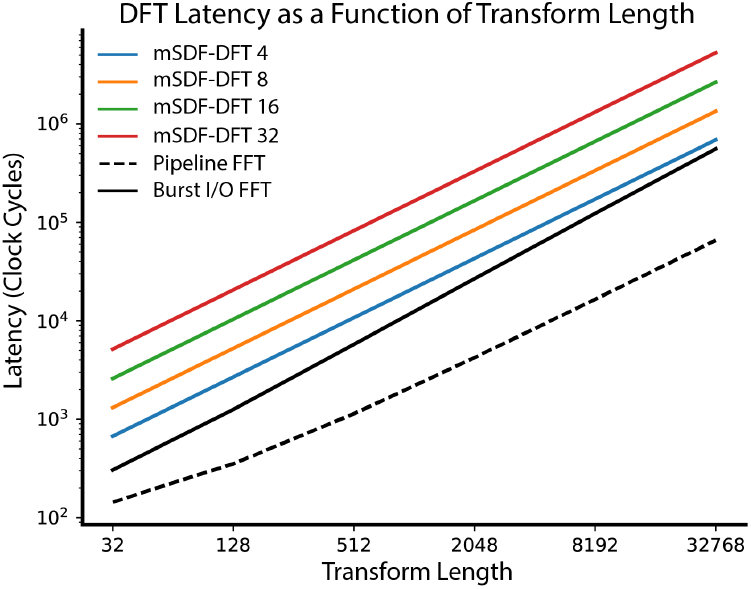
Characterization of latency between mSDF-DFT andFFT methods. Results show that mSDF-DFT throughput is less than FFT methods as expected. However, the choice of mSDF-DFT parameters can bring time performance to burst-I/O FFT levels.

Thus, assuming a constant bandwidth, the latency of the mSDF-DFT is much lower with smaller frame size.

### C. Evaluation of Online Neural Sensing

Finally, we evaluated the mSDF-DFT architecture performance and accuracy on LFPs recorded from an optical thalamocortical deep brain stimulation application. Data was obtained from infrared stimulation of auditory thalamus with recordings made from microwire recordings of auditory cortex in the chronically implanted rat (Figure 5A). Stimulation of the ventral division of auditory thalamus drives excitatory responses in layers III/IV of auditory cortex through a single synapse (Figure 5B). Local field potential responses through this circuit have been extensively characterized, creating an opportune circuit by which to assess mSDF-DFT performance. A characteristic LFP driven by 3 mJ per pulse INS is shown in Figure 5C.

**Fig. 5.**
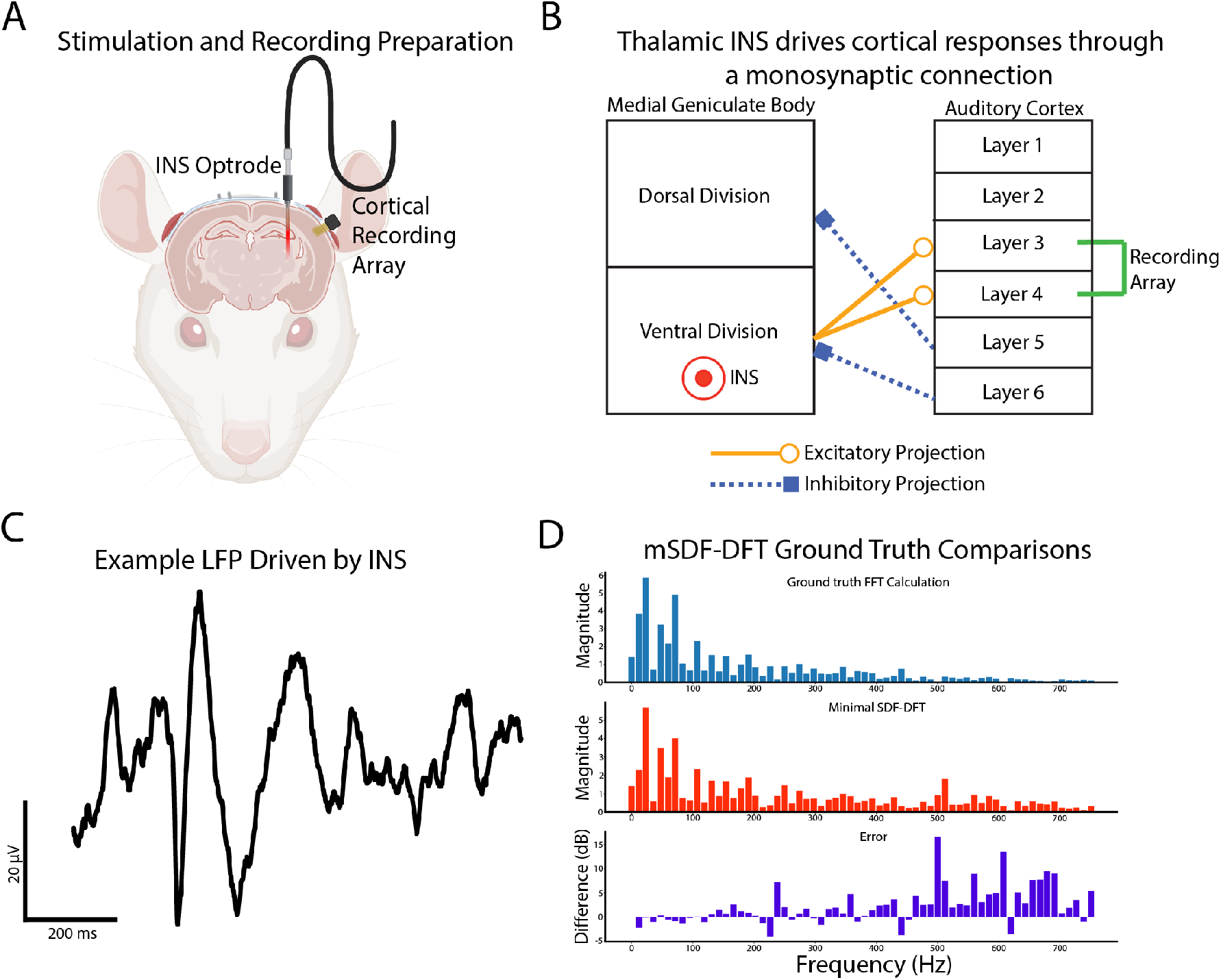
Application of the mSDF-DFT to closed-loop neural sensing. A,B. Recordings were obtained from a chronically implanted rat infrared neural stimulation (INS) preparation. Stimulating optical fibers were placed into auditory thalamus with recordings made from 16 channel microwire arrays implanted in auditory cortex. Thalamic afferents occur across a single synapse to cortex and thus represent direct thalamocortical entrainment from INS. A portion of the figure was constructed in BioRender (www.biorender.com) software. C. Example LFP waveform and time-frequency decomposition showing *β* and *γ* power vs time. D. Example histogram comparison of SDF-DFT and offline FFT power calculations show minimal error in calculation of *β* and *γ* power.

The mSDF-DFT architecture demonstrated superior resource and power efficiency when compared to benchmark architectures (Table 5) consistent with our observed power and resource utilization findings (Figures 2,3). A 33% dynamic power reduced and 4% resource utilization reduction over burst-I/O FFT was observed. A 38% dynamic power reduced and 4% resource utilization reduction over Goertzel Algorithm was observed. A further 85% power reduction and 73% resource utilization reduction over Pipelined FFT was observed.

**TABLE IV.**
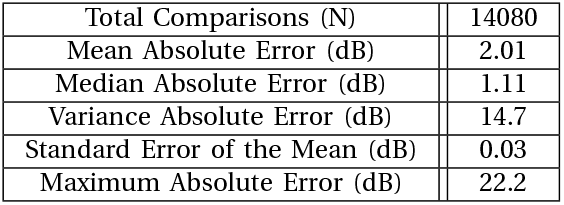
Summary statistics for LFP mSDF-DFT and Benchmark FFT.

|  |  |
| --- | --- |
| Total Comparisons (N) | 14080 |
| Mean Absolute Error (dB) | 2.01 |
| Median Absolute Error (dB) | 1.11 |
| Variance Absolute Error (dB) | 14.7 |
| Standard Error of the Mean (dB) | 0.03 |
| Maximum Absolute Error (dB) | 22.2 |

**TABLE V.** Summary of mSDF-DFT Performance Against Benchmark architectures in Neural Sensing.

|  | mSDF-DFT | Burst I/O FFT | Goertzel Filter | Pipelined FFT |
| --- | --- | --- | --- | --- |
| LUT | 280 | 291 | 722 | 1169 |
| FF | 339 | 688 | 440 | 21941 |
| BRAM | 1 | 1 | 0 | 0 |
| DSP | 2 | 2 | 1 | 9 |
| Resource Utilization | 2.4 | 2.5 | 2.5 | 9.0 |
| Dynamic Power (mW) | 8 | 12 | 13 | 52 |

To evaluate mSDF-DFT’s calculation accuracy, a total of 10, 1 second LFP recordings were utilized. For each sample, eleven 128-point DFT were performed with no overlap, resulting in a total of N=14,080 DFT calculations utilized for accuracy analyses. A characteristic mSDF-DFT and ground truth FFT spectrums and spectral differences is shown in Figure 5D. D’Agostino and Pearson tests for normality[37] show that errors did not fit normal distributions (p<0.05), necessitating the use of mean and median measurements of absolute error as optimal summary statistics for mSDF-DFT error analysis[38]. Error analysis results are summarized in Table 4. The mean absolute error between measured mSDF-DFT and benchmark offline FFT was 2.01 dB with a standard error of the mean of 0.03 dB. The median of absolute error was found to be 1.11 dB, and the absolute maximum error was 22.2 dB. The relatively small mean absolute error suggests that the mSDF-DFT provides accurate low power and resource estimation of online neural signals. Sources of error are more concentrated in higher frequency components and likely due to propagation of memory limitations and quantization errors inherent to online systems at higher frequencies, which can be mitigated by increasing word length[39].

## V. Discussion

We describe the design and implementation of a new and novel power and resource efficient DFT architecutre called the mSDF-DFT. The architecture implements the DFT in a completely serialized fashion with inverse transform capability that possesses all the features available in SOTA Xilinx FFT IP cores. It is important to note that our data does show well-defined use cases for the mSDF-DFT. When minimization of power is the desired constraint, the mSDF-DFT is superior in applications with moderate bin counts and transform lengths and completely outperforms pipelined FFTs in power performance. Similarly, at moderate *N*_*B*_ to N given by Equation 7, the mSDF-DFT can outperform burst I/O FFT methods. Furthermore, the mSDF-DFT almost completely outperforms Goertzel filter in dynamic power, except at extremely small *N*_*B*_ to N ratio, where Goertzel filter’s efficient memory access pattern is salient. Therefore, the area of best power performance can be characterized as a moderate *N*_*B*_ to N ratio, which is precisely where neural sensing, among other medical devices, operates. Thus, mSDF-DFT offers high-resolution spectral decomposition with minimal power consumption in any application where power constraints represent critical design parameters.

We also find use cases where the mSDF-DFT provides minimal resource usage. Although this metric (Equation 4) provides a holistic estimate of total system use, resource usage is defined as percentage utilization and is therefore FPGA-dependent. Specifically, the different component categories, i.e., DSP, LUT, etc., are weighted inversely to their quantity present in the FPGA. Although the resulting percentages presented in resource consumption may not transfer to FPGA architectures with significantly different component composition, the relative absolute utilization should be conserved.

### A. Considerations for Energy Efficient DFTs

While throughput for the mSDF-DFT is lower than massively parallelized methods such as pipelined FFT implementations, most power constrained applications aim to minimize with neural sensing and control systems. This is done by having sampling and throughput rates that adequately capture neural responses but at much lower speeds than typical FPGA clock speeds[40]. In a system that involves real-time signal acquisition and concurrent transform into the frequency domain, the maximum number of frequency bins that can be processed within the timeframe is given by:

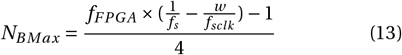

where *f*_*FPGA*_ is the FPGA clock frequency, *f*_*s*_ is the sampling frequency, *f*_*sclk*_ is the SPI frequency and *w* is the bit depth of the SPI transmission. The term 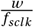 compensates for the SPI transmission time. Neural sensing typically involves a low sampling rate, e.g. 1 kHz for LFP recording, 100 Hz for intracranial EEG systems, and minimum 7 kHz for spike detection[41], [42]. At this sampling rate, assuming 50 MHz FPGA clock frequency and 1 MHz SPI frequency, 1600 to 124,000 frequency bins can be processed with the proposed mSDF-DFT, which is significantly above minimal requirements for online neural sensing and measurement applications.

As implemented, the mSDF-DFT provides an accurate and low-resource usage DFT implementation. However, we do show that some small approximation error does exist in mSDF-DFT estimation, which may compound if bin size and transform lengths grow large. It is possible to further reduce round off and propagation errors by implementing fault tolerance methods, such as online memory[43] or algorithm based[44] fault tolerance into our mSDF-DFT architecture. However, fault correction methods will likely add to power and resource consumption, necessitating careful optimization of the accuracy-resource tradeoff. It was also observed that mSDF-DFTs are particularly poised for use in low to moderate transform length and bin size applications. We believe that more power and resource savings can be achieved with direct implementation of mSDF-DFT into application specific integrated circuits (ASICs) which would allow for full optimization of power consumption using only necessary resources providing best performance. Lastly, it is important to note that the mSDF-DFT does not only outperform Goertzel filter in power and resource utilization. It is well known that Goertzel filters exhibit numerical instability for fixed point arithmetic and long input sequences owing to its use of purely real digital filters for Fourier coefficient calculation[45]. This can be easily avoided by using true DFT estimation of Fourier coefficients. Thus, the mSDF-DFT represents a promising high-performance alternative to the Goertzel filter. While the mSDF-DFT was tested on Xilinx FGPA platforms, our results represent physical measurements of gate utilization and power consumption, meaning other FPGA platforms will see similar reductions in power and resource utilization. Furthermore, the defined mSDF-DFT architecture is immediately scalable to implementation on ASIC designs, and is intended for future work.

### B. Extended Applications of the mSDF-DFT

While the proposed DFT is of interest for current clinical implementations of adaptive DBS, we envision this architecture able to facilitate the investigation of more complex control methods in using online encoding and decoding[46], [47], brain-machine interfaces utilizing spectral decomposition methods[48]–[50], or as a plug-in tool for other neural sensing and recording platforms necessitating online spectral estimation and/or closed-loop control[51], [52]. This architecture also extends generally to non-medical applications such as space systems and satellite instrumentation[53]–[55] and wireless communication systems[56], [57], or in any application where conserved power and resource usage is desired.

A particularly promising application of the mSDF-DFT architecture is its integration into Fourier transform-enabled deep learning accelerators[58], [59]. Deep neural network model inference using Fourier computations is becoming increasingly popular in embedded deep learning, especially in real-time signal and image processing applications[60], [61] using traditional FFTs for feature extraction, conversion of convolution operations into simpler multiplications, and pattern matching and detection. However, conventional pipeline and burst-mode FFT architectures have substantial energy consumption and impose considerable demands on silicon wafer area. In contrast, mSDF-DFTs are well-suited for on-chip deep learning applications and can serve as efficient components within neural network accelerators due to their low gate count and minimal resource utilization. This efficiency enables either a reduction in overall die size or the reallocation of chip area to other critical functions. Moreover, the low implementation overhead of mSDF-DFTs allows for parallel deployment with configurable bin sizes tailored to extract relevant features, facilitating the development of power- and resource-efficient hardware DFT accelerators.

## VI. Conclusion

We show that the defined mSDF-DFT architecture outperforms low-power gold standard benchmark Goertzel algorithm DFTs and burst I/O FFTs with a well-defined use cases in biopotential sensing. Specifically, we observe a 33% reduction in dynamic power and 4% reduction in resource utilization in a neural sensing application when compared to SOTA burst I/O FFT. The mSDF-DFT has greater latency when compared to benchmark FFTs but achieves high accuracy transforms at state-of-the-art low-power consumption, making the mSDF-DFT a potent tool for neural sensing applications and beyond.

## Code and Data Availability

Raw and processed LFP data can be found at the following Open Science Framework data repository: https://osf.io/fb48z/.

## Disclosures

R.Y., K.A.L, and B.S.C hold a provisional patent on the technology described in this manuscript. K.A.L. is a co-founder and equity holder for Neuronoff, Inc. K.A.L. is also a co-founder and equity holder of NeuraWorx. K.A.L. is a scientific board member and has stock interests in NeuroOne Medical Inc. K.A.L. is also a paid member of the scientific advisory board of Abbott and Presidio Medical, and a paid consultant for the HuMANNity, ONWARD and Restora Medical. H.D.O holds patents related to low power FFT implementations. H.D.O is also a consultant for Inspire Medical Systems. B.S.C is an unpaid scientific consultant for BECATech Inc.

## Acknowledgments

The authors would like to thank the following for helpful feedback for this manuscript: James Trevathan PhD, Suyash Bhatt PhD, and Claudia Krogmeier PhD. This study was supported by grants from the National Institutes of Health (NINDS #RF1-NS129955, PI: K.A.L.) and the Hilldale Undergraduate/Faculty Research Fellowship (University of Wisconsin-Madison, R.Y.)

**Richard Yang** is an undergraduate research assistant at the Wisconsin Institute for Translational Neuroengineering. He is currently pursuing a bachelor’s degree in biomedical engineering and computer science at the University of Wisconsin-Madison.

**Heather D. Orser** received her BSEE from Minnesota State University, Mankato and her MSEE and PhD from the University of Minnesota. She is currently an assistant professor of Electrical and Computer Engineering at the University of St Thomas, St. Paul, MN.

Prior to her time at St Thomas, Heather worked in the development of implantable neuromodulation systems at both Inspire Medical and Medtronic where she led the development of a number of next-generation systems and successfully assessed the safety of implantable devices for patients undergoing MRIs. Her research focuses on the development of neuromodulation systems for use in research and the clinic.

**Kip A. Ludwig** received his bachelor’s degree in Biomedical Engineering from Arizona State University, and his Masters and PhD in Biomedical Engineering from the University of Michigan. He is currently an Associate Professor in the Departments of Neurological Surgery and Surgery at the University of Wisconsin-Madison, and the Co-Director for the Wisconsin Institute for Translational Neuroengineering (WITNe).

Prior to his time at UW-Madison, Dr. Ludwig worked in both industry and government. While at CVRx® he led the development of the ‘Neo’ electrode to treat hypertension and heart failure which has been subsequently FDA PMA approved. He also co-led the translational devices program at the National Institute for Neurological Disorders and Stroke, and led the trans-NIH translational neurotechology programs under the NIH SPARC and BRAIN Initiatives. His research focuses on accelerating the path to translation for next-generation devices to hack the nervous system to treat a variety of diseases/disorders inadequately managed by drugs/biologic.

**Brandon S. Coventry** received a bachelor’s degree in electrical engineering from Saint Louis University, a master’s degree in electrical and computer engineering from Purdue University, and a PhD in biomedical engineering from Purdue University. He is currently a postdoctoral research associate at the University of Wisconsin-Madison, WI USA in the Department of Neurological Surgery and the Wisconsin Institute for Translational Neuroengineering. His research interests include design of next generation implantable pulse generators, mechanisms of deep brain stimulation, translation of neurotechnologies, and thalamic neuroscience.

## Notes

This work was supported by the United States National Institute of Neurological Disorders and Stroke (NINDS) grant #RF1-NS129955 and the Hilldale Undergraduate/Faculty Research Fellowship (University of Wisconsin-Madison).

https://osf.io/fb48z/

### Summary of Updates

Revised version after first round of review with updates to IEEE membership status.

